# Novel BRAF fusions in pediatric histiocytic neoplasms define distinct therapeutic responsiveness to RAF paradox breakers

**DOI:** 10.1101/2020.04.13.039032

**Authors:** Payal Jain, Lea F. Surrey, Joshua Straka, Pierre Russo, Richard Womer, Marilyn M. Li, Phillip B. Storm, Angela Waanders, Michael D. Hogarty, Adam Resnick, Jennifer Picarsic

## Abstract

Pediatric histiocytic neoplasms are clonal hematopoietic disorders driven by mutations activating the mitogen-activated protein kinase (MAPK) pathway, such as BRAF-V600E. In non-BRAFV600E cases, we investigated alternative MAPK mutations and found two novel BRAF gene fusions. We investigated the distinct responsiveness of novel BRAF fusions to RAFi therapies and explored the mechanistic basis of such differential responses compared to other BRAF fusions. Two histiocytic patient tumors were analyzed using the CHOP Comprehensive Next-Gen Sequencing Solid Tumor Panel and a targeted RNA-seq panel for 106 fusion partner genes. In the two M- and L-type histiocytic neoplasms assessed, we found novel and rare BRAF gene fusions, MTAP-BRAF and MS4A6A-BRAF, respectively. Both BRAF fusions activated the MAPK/ PI3K pathways and showed homo- and hetero-dimerization with BRAF and the respective N-terminal fusion partner. In contrast to common BRAF fusions, MTAP-BRAF and MS4A6A-BRAF did not respond to PLX8394 due to a lack of disruption of active fusion homo- and hetero-dimers, which was in turn due to the untargeted, stable dimerization mediated by the N-terminal fusion partners. Conversely, we observed robust suppression with LY3009120 that bound fusion dimers and kept them in an inactivate confirmation. MEKi were found to successfully suppress fusion driven signaling and oncogenic phenotypes. Our finding that PLX8394 does not disrupt MTAP-BRAF or MS4A6A-BRAF dimerization due to contribution of N-terminal partners defines a novel paradigm for the distinct mechanisms sought by BRAF fusions in response to RAFi therapy. Overall, this study highlights the unique and differential biology hijacked by BRAF fusions in response to RAFi and further warrants detailed mechanistic classification of BRAF fusions based on their responsiveness to targeted agents.

---

Histiocytic neoplasms are a diverse group of clonal hematopoietic disorders that are driven by mutations activating the mitogen-activated protein kinase (MAPK) and phosphoinositide 3-kinases (PI3K) pathways^1–3^. While BRAF-V600E is the most common alteration in histiocytic neoplasms, multiple alternate pathway activating mechanisms have been described, including *MAP2K1*, *ARAF*, *PIK3CA*, *NRAS*, and *KRAS* mutations as well as *BRAF*, *ALK*, and *NTRK1* fusions^1–3^. BRAF-fusions previously reported in cases of histiocytic neoplasms^1,4–7^ were found to contain the N-terminal region of another gene (often of unclear significance) joined to the BRAF kinase domain (including exons 9-18 or 11-18), resulting in the loss of the BRAF N-terminal regulatory RAS-binding regions (exons 1-8). Despite the prevalence of BRAF-alterations in histiocytic tumors, to date there have been no detailed molecular investigations comparing BRAF-fusions found in distinct sub-types of histiocytic neoplasms and only one study explored responsiveness of such BRAF-fusions to single-agent RAF-therapies^4^. To address this, we present two pediatric histiocytosis cases with divergent pathologic and clinical features, each harboring a novel BRAF-fusion as identified by a clinically validated next-generation sequencing panel (Supplemental Methods and Table 1) and study their *in vitro* responsiveness to RAF-targeted inhibitors.

Case 1, a 16 year-old female with a 2.5 cm rapidly growing subcutaneous thigh mass was diagnosed with a malignant histiocytic neoplasm (“M group”)^8^, with a phenotype spanning histiocytic sarcoma (CD163/CD14/CD68+) and Langerhans cell sarcoma (CD1a/Langerin/S100) with a modestly elevated Ki-67 proliferation index (up to 20%) (Figure 1A-F). Targeted RNA-sequencing identified a fusion of *MTAP* (NM_002451.3) exons 1-7 to *BRAF* (NM_004333.4) exons 9-18. Resection margins were negative. The patient is disease-free three years post-resection. Case 2, a 12-year-old female with a 5.3 cm rapidly enlarging heel mass encasing and invading the calcaneus was diagnosed with a juvenile xanthogranuloma (JXG) family lesion (CD163/CD68/CD14/fascin/Factor XIIIa+) (Figure 1G-M). Despite the lack of cytologic atypia or increased mitotic rate by H&E stain, a Ki-67 proliferation index stain revealed increased staining in lesional cells up to 40%. Targeted RNA-seq identified a fusion of *MS4A6A* (NM_022349.3) exons 1-6 and *BRAF* (NM_004333.4) exons 11-18. During staging, the patient was found to have PET-avid dissemination to lymph nodes and lung (Figure 1N-P). While the morphologic features were consistent with a low-grade histiocytic lesion of JXG phenotype, the integration of the 1) high Ki-67 proliferation index, 2) aggressive clinical behavior with lymphatic/metastatic-like spread, and 3) novel molecular *BRAF-*fusion suggested an atypical JXG family neoplasm with uncertain biological behavior. The patient was treated with 12 cycles of clofarabine with clinical remission of metastatic sites and near clinical remission at primary site now 12 months off therapy.

**Figure 1.**
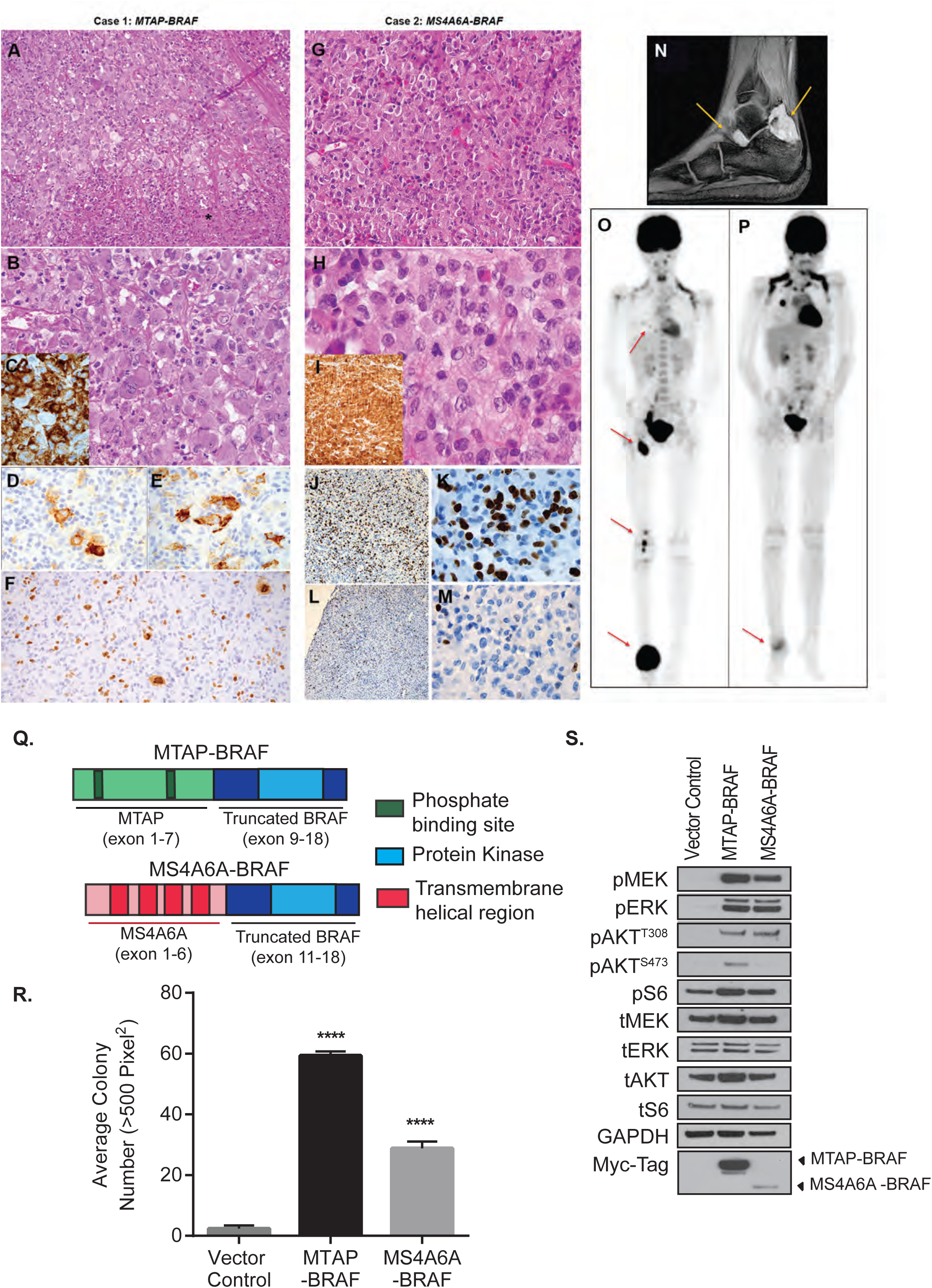
Novel BRAF-fusions in histiocytic neoplasms mediate oncogenicity via activation of MAPK/PI3K/mTOR pathway: Malignant histiocytic neoplasm with histiocytic and Langerhans cell sarcoma phenotypes with novel *MTAP-BRAF* fusion, and atypical juvenile xanthogranuloma family lesion with novel *MS4A6A-BRAF* fusion. **A-F.** Case 1 Malignant histiocytic neoplasm with large, pleomorphic cells (A-B) and areas of necrosis (*). Immunohistochemistry with CD163 (C), CD1a (D) and Langerin (E) in a subset of lesional cells. Ki-67 proliferation index (F) was elevated up to 20%, including atypical large cells (F). (Original magnification A. 200x, B, 4000x, C-E. 1000x, F. 200x). BRAF VE1 immunostain was negative (not shown). **G-M.** Case 2 atypical juvenile xanthogranuloma (JXG) family neoplasm with bland histiocytes (G-H) and a rare mitosis (H, center). Immunohistochemistry with Factor XIIIa (I) was strongly and diffusely positive. The Ki-67 proliferation index was variable, as high as 40% (J-K) in one core biopsy and as low as 10% in other core (L-M) taken at the same time and accounting for inflammation, which was low in both core biopsies. (Original magnification: G. 100x, H. 1000x, I. 200x, J. 100x, K. 1000x, L. 100x, M. 1000x). The BRAF VE1 immunostain was negative (not shown). **N-P.** Case 2 with JXG: Imaging at diagnosis revealed a crescentic enhancing soft tissue mass by magnetic resonance imaging wrapping around the calcaneus, deep to the Achilles tendon (N, arrows) and positron emission tomography (PET) scanning revealed abnormal signal in the ankle (primary), knee, inguinal region and chest (O). Following 9 of 12 cycles of clofarabine, PET scan revealed resolution of disseminated disease and shrinkage of the primary ankle tumor. **Q**. Structure of novel BRAF-fusions in histiocytic neoplasms. MTAP-BRAF: MTAP exons 1-7 encode phosphate binding sites, trimerization site at Trp189 residue, and substrate binding site, and BRAF exons 9-18 encode the tyrosine kinase domain. MS4A6A-BRAF: MS4A6A exons 1-6 encode 4 transmembrane helical regions, and BRAF exons 11-18 encode the tyrosine kinase domain. **R.** Soft agar assay using NIH3T3 cells stably expressing BRAF-fusions. Error bars represent SEM, n=5, ***p-value<0.001 compared with control conditions. **S.** Western blot analysis of MAPK and PI3K/mTOR pathway proteins in NIH3T3 cells stably expressing BRAF-fusions. ‘p-‘ and ‘t-‘ represent phosphorylated and total versions of protein, respectively.

MTAP-BRAF and MS4A6A-BRAF fusions are predicted to contain all functional domains of MTAP and MS4A6A, respectively, along with the BRAF kinase domain but no N-terminal regulatory, RAS-binding domain (Figure 1Q). For molecular and therapeutic characterization, *MTAP-BRAF* and *MS4A6A-BRAF* were cloned and stably expressed in a heterologous cell model since patient-derived cell lines were lacking. The NIH/3T3 cells model system was utilized due to its ability to reliably discern oncogenic fusion profiles^9–11^. In soft agar assays, both *MTAP-BRAF* and *MS4A6A-BRAF* expressing NIH3T3 showed a significant increase in colony count over control (p<0.0001, Figure 1R). Next, we tested activation of downstream MAPK and PI3K/mTOR pathways. Under serum starved conditions, we observed elevated levels of phosphorylated-ERK and -S6 in both BRAF-fusion expressing cells compared to controls, indicating aberrant activation of both the MAPK and PI3K/mTOR pathway, respectively (Figure 1S). Slightly higher PI3K/mTOR pathway activation levels in MTAP-BRAF versus MS4A6A-BRAF cells could be partly explained by higher MTAP-BRAF protein expression (Figure 1S, Myc-tag blot).

A single report on BRAF-fusions in LCH^4^ has shown unresponsiveness to BRAF-V600E specific inhibitors such as vemurafenib, termed first-generation RAF inhibitors (RAFi), but observed suppression by second-generation RAFi, PLX8394, and downstream MEK inhibition, similar to other pediatric glioma studies on BRAF-fusions^9,11^. Herein, we evaluated the responsiveness of novel MTAP-BRAF and MS4A6A-BRAF to such targeted inhibitors. Upon targeting the NIH3T3 models with first-generation RAFi PLX4720, as expected, no suppression of BRAF-fusion driven signaling or growth was observed (Supplemental Figure 1A). Interestingly, second-generation RAFi PLX8394 also showed no suppression in MTAP- or MS4A6A-BRAF driven soft agar growth despite targeting MAPK/PI3K signaling (Figure 2A-B). <u>This</u> is in contrast to PLX8394-mediated suppression of BRAF-fusion driven growth in the previously described LCH^4^ and other cancers, such as the KIAA1549-BRAF fusion in pediatric glioma^9–11^. PLX8394 suppressed FAM131B-BRAF (a pediatric glioma derived fusion^12,13^) and BRAF-V600E driven growth and signaling as well as actively disrupted FAM131B-BRAF dimers (Supplemental Figures 1B-D), highlighting therapeutic differences between MTAP-/MS4A6A-BRAF, BRAF-V600E and other BRAF-fusions.

**Figure 2.**
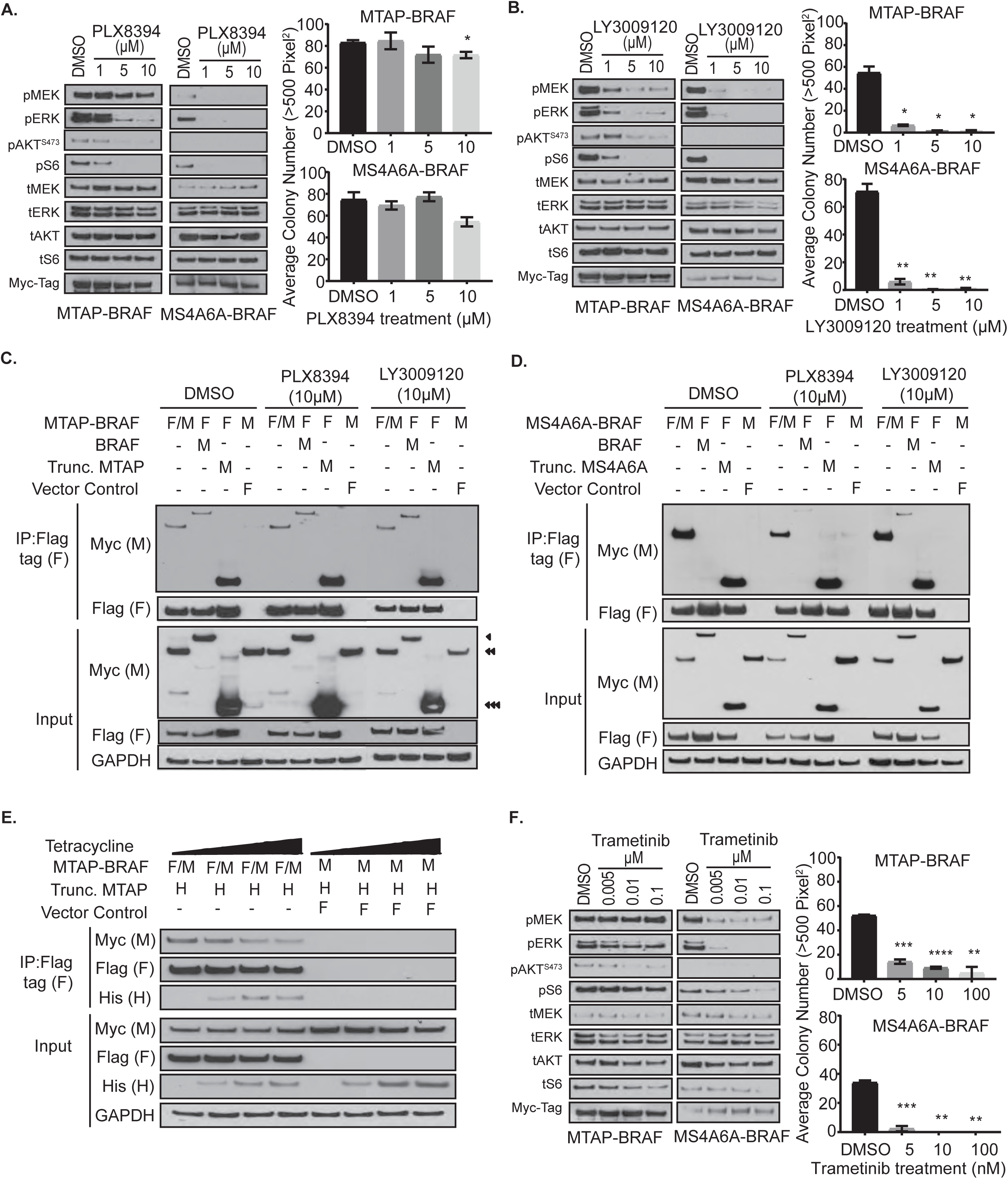
MTAP-BRAF and MS4A6A-BRAF fusions are not suppressed by second generation RAF inhibitors but demonstrate sensitivity to LY3009120 and MEK inhibitors. **A.** Western blot analysis (left) and soft agar colony counts (right) showing the effect of second generation RAFi, PLX8394, on NIH3T3 cells expressing MTAP-BRAF and MS4A6A-BRAF respectively. **B.** Western blot analysis (left) and soft agar colony counts (right) showing the effect of pan-RAF-dimer inhibitor, LY3009120, on NIH3T3 cells expressing MTAP-BRAF and MS4A6A-BRAF respectively. **C.** Co-immunoprecipitation (co-IP) assay assessing homo-dimerization of MTAP-BRAF as well as hetero-dimerization with wild-type BRAF and Trunc-MTAP in HEK293 cells under control, PLX8394, and LY3009120 treated conditions. **D.** Co-immunoprecipitation assay assessing homo-dimerization of MS4A6A-BRAF as well as hetero-dimerization with wild-type BRAF and Trunc-MS4A6A in HEK293 cells under control, PLX8394, and LY3009120 treated conditions. **E.** Competition co-IP assay assessing preferential interaction of Trunc. MTAP with MTAP-BRAF fusion versus homo-dimerization. Increasing doses of tetracycline (0, 0.1, 0.5, 1 ug/ml) used to regulate protein level of His-tagged Trunc-MTAP. **F.** Western blot analysis (left) and soft agar colony counts (right) showing the effect of MEK inhibitor, trametinib, on NIH3T3 cells expressing MTAP-BRAF and MS4A6A-BRAF respectively. Error bars represent SEM, n=3. No value on bar represents NS (non-significant), *p-value<0.05, **p-value< 0.01, ***p-value<0.001 compared with control conditions. ‘p-‘ and ‘t-‘represent phosphorylated and total versions of protein, respectively.

BRAF-fusions function as active homo- and heterodimers (with wild-type BRAF) to mediate cell signaling^9,11^. We found that MTAP- and MS4A6A-BRAF also mediate such protein-protein interactions in co-immunoprecipitation assays (Figure 2C-D, DMSO lanes). PLX8394 blocks BRAF kinase activity via disrupting BRAF dimerization^14^ but we observed no disruption of MTAP- and MS4A6A-BRAF fusion dimerization with PLX8394 (Figure 2C-D, PLX8394 lanes), thereby providing a plausible explanation for PLX8394 unresponsiveness in soft agar assays though MAPK/PI3K signaling remains discordantly suppressed by some unknown mechanism. This distinct unresponsiveness to pan-RAFi represents a significant departure from the current view that *BRAF*-fusions and other *BRAF* mutations should respond to second-generation RAFi such as PLX8394^9,15^. We found that this difference arises due to the contribution of N-terminal partners, MTAP (exons 1-7) and MS4A6A (exons 1-6), to respective fusion dimerization that is unaffected by PLX8394 (Figure 2C-D, lanes 3,7). Similar role of N-terminal partner accounts for differential response of *CRAF-*fusions to PLX8394^10^. Furthermore, we observed that Trunc-MTAP (exons 1-7) competitively substituted MTAP-BRAF homo-dimerization in a dose-dependent manner, suggesting preferential and potent protein interactions mediated by N-terminal partner in these histiocytic-specific BRAF-fusions (Figure 2E).

To target dimerization-dependent oncogenicity of MTAP- and MS4A6A-BRAF via a different mechanism, we tested LY3009120, a pan-RAF dimer inhibitor that binds and stabilizes the BRAF dimer in an inactive conformation^16^. LY3009120 showed robust suppression of both fusion-mediated signaling and colony transformation (Figure 2B) while stabilizing the MTAP- or MS4A6A-BRAF in inactive conformation (Figure 2C-D, respectively, <u>lanes 9-11</u>). We also tested the effect of FDA-approved MEK inhibitors (MEKi)^17^, selumetinib and trametinib. We observed dose-dependent decrease in phospho-ERK and growth with trametinib (Figure 2F) and selumetinib (Supplemental Figure 2) in both BRAF-fusion models suggesting downstream MEKi as a therapeutic alternative to RAFi.

Though functional data predicts similar effects for these two novel fusions, each neoplasm had discordant clinical and pathologic phenotypes. Malignant histiocytic lesions typically have aggressive behavior^18^, unlike case 1. Typical JXG family lesions often show indolent behavior in pediatric patients^19^. Even in rare systemic presentations, they do not feature a lymphatic-type dissemination, as demonstrated in case 2. Furthermore, Ki-67 proliferation index in JXG lesions is typically less than 20%, and no higher than 10% (unpublished data, JP). Thus, the focally high proliferation rate (40%) in the *MS4A6A-BRAF* JXG family lesion was the first correlate to its aggressive clinical behavior. On the contrary, the *MTAP-BRAF* malignant histiocytic neoplasm had only modestly elevated proliferation rate (up to 20%), which is lower than most malignant histiocytic neoplasms (often >30%)^20^. This may correlate with its more indolent behavior, despite its high-grade cytologic features. Both of these unusual, divergent phenotypes further emphasize that in histiocytic neoplasms, an integrated diagnosis with pathologic, molecular, and clinical/radiographic correlation are all needed for best diagnosis and allows for expanded treatment options.

## Acknowledgments

The authors would like to thank Dr. Ronald Jaffe for second review of the cases and appraisal of the manuscript.

This work was supported by the Children’s Brain Tumor Tissue Consortium funding sources (P.J., A.J.W., P.B.J., and A.C.R).

## Authorship Contribution

J.P., L.F.S, P.J. conceived the study; J.P., L.F.S and P.R. generated, analyzed and interpreted the pathology data; P.J. and J.S. performed molecular experiments as well as analyzed and interpreted the results; P.J., L.F.S, J.S. and J.P. wrote the manuscript and compiled the figures; P.J., L.F.S, J.S., P.R., R.W., M.L., P.B.S., A.J.W., M.D.H., A.C.R and J.P. edited and reviewed the manuscript.

