## Supplemental methods, table and figures. for "Novel BRAF fusions in pediatric histiocytic neoplasms define distinct therapeutic responsiveness to RAF paradox breakers"

**Supplemental Material:**

**Supplemental Figure 1: Novel BRAF fusions are not suppressed by first generation RAFi, and BRAF-V600E and FAM131B-BRAF are suppressed by both second generation RAFi and LY3009120.**

(A) Western blot analysis (top) and soft agar colony counts (bottom) showing the effect of 1st (PLX4720) generation RAFi on NIH3T3 cells expressing MS4A6A-BRAF and MTAP-BRAF respectively. (B) Western blot analysis (top) and soft agar colony counts (bottom) showing the effect of second generation RAFi, PLX8394 (left panel) and LY3009120 (right panel) on NIH3T3 cells expressing mutant BRAF-V600E. (C) Western blot analysis (top) and soft agar colony counts (bottom) showing the effect of second generation RAFi, PLX8394 (left panel) and LY3009120 (right panel) on NIH3T3 cells expressing FAM131B-BRAF fusion. (D) Co-immunoprecipitation assay assessing homo-dimerization of FAM131B-BRAF as well as hetero-dimerization with wild-type BRAF in HEK293 cells under control, PLX8394, and LY3009120 treated conditions. Error bars represent SEM, n=3. No value on bar represents NS (non-significant), *p-value<0.05, **p-value< 0.01, ***p-value<0.001 compared with control conditions.

**Supplemental Figure 2: Novel BRAF fusions are targeted by MEK inhibitors.** Western blot analysis (top) and soft agar colony counts (bottom) showing the effect of MEK inhibitor, selumetinib, on NIH3T3 cells expressing MS4A6A-BRAF and MTAP-BRAF respectively.

**Supplemental Table 1: Results of Comprehensive Solid Tumor Panel showing RNA based fusion results and DNA sequence results from 238 genes. Abbreviations: VAF = variant allele fraction**

**Supplemental Methods:**

**Histologic evaluation and Next Generation Sequencing:**

Cases were ascertained from the files of the Children's Hospital of Philadelphia (CHOP) and reviewed in consultation at the UPMC Children's Hospital of Pittsburgh. Histologic evaluation and immunohistochemistry was performed including CD1a (EP3622, Cell Marque), CD3 (Polyclonal, Dako), CD14 (EPR3653, Cell Marque), CD68 (KP1, Dako), CD163 (10D6, Leica), S100 (Polyclonal, Leica), Langerin (CD207, 12D6, Lifespan Biosciences), fascin (55K-2, Dako), Factor XIIIa (N1N3, GeneTex), BRAF V600E (VE1, Ventana Medical System), and Ki-67 (MIB-1, Dako).

Neoplasms were analyzed on the CHOP Comprehensive Next Generation Sequencing Solid Tumor Panel, as previously described[1]. This panel includes tumor-only RNA-seq by anchored multiplex PCR for 110 fusion partner genes and DNA sequencing of 238 genes. Fusion genes were confirmed by standard reverse transcription and Sanger sequencing.

**Fusion cloning and characterization:**

MTAP-BRAF and MS4A6A-BRAF were cloned and transduced into heterologous NIH3T3 cells for stable expression of BRAF-fusions, as previously described[2]. Signaling pathways activated by fusions were analyzed in serum starved conditions via western blotting for phosphorylated proteins. The antibodies used, pMEK (#9154), tMEK (#4694), pERK (#4370), tERK (#4695), pAKT Thr308 (#4056), pAKT Ser473 (#9271), tAKT (#2920), pS6 (#4858), and tS6(#2317), were purchased from Cell Signaling (1:1000). Oncogenicity was assessed using soft agar transformation assays[2]. For creation of a Tet-inducible plasmid, we used pT-REx-DEST Gateway™ Vectors with a His-tag and Tetracycline inducer catalog # A39246 (Life Technologies) and for protein detection, we used His-Tag (D3I1O) XP® Rabbit mAb ( #12698).

**Cellular Drug Assays**

Selumetinib (Astra Zeneca, Gaithersburg, MD, USA) and LY3009120 (Eli Lily, Indianapolis, IN, USA) were purchased through Selleckchem (Houston, TX, USA). PLX4720 and PLX8394 were provided by Plexxikon (Berkeley, CA), and trametinib was supplied by GlaxoSmithKline. Cells were plated at 1x10^6 cells/well in a 6-well plate and serum starved for 24 hours followed by exposure to specific drug for 1 hr. All drugs were dissolved in dimethyl sulfoxide and kept at -20$^{o}$C.

**Co-immunoprecipitation (Co-IP) assay:**

Interactions of MTAP-BRAF and MS4A6A-BRAF with themselves, with truncated MTAP (exons 1-7) and truncated MS4A6A (exons 1-6), respectively, as well as wild-type BRAF were interrogated through co-transfections of Myc and Flag-tagged constructs into HEK293 cells. Transfections were performed using Lipofectamine 2000. 24 hours after transfection, cells were serum starved for 24 hours. Cells were then treated with indicated drug concentrations for an hour, lysed with RIPA buffer, centrifuged and lysates were normalized. Anti-Flag antibody coated beads (M8823, Millipore-Sigma, Burlington, MA, USA) were utilized to immunoprecipitate the tagged proteins at 4$^{o}$C for 2 hours. Three 15 minute washes were performed at 4$^{o}$C followed by a PBS wash and elution using 2X LDS (Lithium dodecyl sulfate). Samples were heated at 70$^{o}$C for 10 minutes and analyzed through western blotting. Detection of tagged protein expression was accomplished using Anti-Myc HRP Antibody (Invitrogen R951-25, 1:5000) and Anti-Flag HRP antibody (Sigma A8592, 1:5000).

**Competition Co-IP assay:**

For the His-tagged, Tet inducible trunc-MTAP plasmid, we used T-REx™ Cell Lines (Thermo Fischer R71007), which stably express the tetracycline repressor protein. Triple transfection with Myc- and Flag-tagged MTAP-BRAF fusion plasmids, along with His-tagged, Tet inducible trunc-MTAP plasmid was done using Lipofectamine 2000 (Life Technologies). 24 hours after transfection, cells were serum starved for 24 hours and treated with increasing doses of Tetracycline (0, 0.1, 0.5, 1 ug/ml), followed by lysis and co-IP as described above.

**Supplemental Table 1: Results of Comprehensive Solid Tumor Panel showing RNA based fusion results and DNA sequence results from 238 genes. Abbreviations: VAF = variant allele fraction**

**Table 1a. Alterations Identified**

| Case # | Fusion Alteration | Tier 3 Variants of Uncertain significance Identified in Sample |
|---|---|---|
| 1 | *MTAP* (NM_002451.3) exon 7 - *BRAF* (NM_004333.4) exon 9 | *FGFR4* (NM_213647.2) c.502G>A (p.V168I), rs150920445, VAF = 0.40<br>*JAK2* (NM_00497203) c.368G>T (p.W123L), VAF = 0.06<br>*NOTCH1* (NM_017617.3) c.4031C>T (p.T1244M), rs201215245, VAF = 0.45<br>*IGF1R* (NM_000875.4) c.1849G>A (p.V617I), VAF = 0.42<br>*TSC2* (NM_00548.3) c.2887G>A (p.V963M), rs45517275, VAF = 0.39 |
| 2 | *MS4A6A* (NM_022349.3) exon 6 - *BRAF* (NM_004333.4) exon 11 | *SPEN* (NM_015001.2) c.9735_9740delCACCC (p.Thr3246_Pro3247del), rs773711901, VAF = 0.33<br>*STAG2* (NM_006603.4) c.3334A>G (p.Thr1112Ala), VAF = 0.41<br>*TSC2* (NM_00548.4) c.4142C>A (p.Pro1381His), rs39515264, VAF = 0.46<br>*CTNNB1* (NM_001904.3) C.1655C>A (p.Ser552Tyr), VAF = 0.46<br>*ARID1B* (NM_020732.3) c.1178C>G (p.Ala393Gly), VAF = 0.53 |

**Table 1b. All genes tested in panel sequencing**

| HGNC-approved symbol | No Clinically Significant Variants or Variants of Uncertain Significance Detected |
|---|---|
| TNFRSF14 | X |
| MTOR | X |
| SPEN | see "Alterations Identified" |
| SDHB | X |
| ARID1A | X |
| MPL | X |
| MUTYH | X |
| CDKN2C | X |
| JUN | X |
| JAK1 | X |
| FUBP1 | X |
| NRAS | X |
| TENT5C | X |
| NOTCH2 | X |
| MCL1 | X |
| NTRK1 | X |
| SDHC | X |
| DDR2 | X |
| CDC73 | X |
| MDM4 | X |
| IKBKE | X |
| H3F3A | X |
| AKT3 | X |
| MYCN | X |
| DNMT3A | X |
| ALK | X |
| MSH2 | X |
| MSH6 | X |
| XPO1 | X |
| ACVR1 | X |
| NFE2L2 | X |
| SF3B1 | X |
| IDH1 | X |
| ERBB4 | X |
| BARD1 | X |
| PDCD1 | X |
| VHL | X |
| RAF1 | X |
| TGFBR2 | X |
| MLH1 | X |
| MYD88 | X |
| CTNNB1 | see "Alterations Identified" |
| SETD2 | X |

| | |
|---|---|
| RHOA | X |
| BAP1 | X |
| PBRM1 | X |
| MITF | X |
| FOXP1 | X |
| EPHA3 | X |
| GSK3B | X |
| GATA2 | X |
| EPHB1 | X |
| FOXL2 | X |
| ATR | X |
| PIK3CA | X |
| SOX2 | X |
| BCL6 | X |
| TP63 | X |
| FGFR3 | X |
| NSD2 | X |
| PHOX2B | X |
| PDGFRA | X |
| KIT | X |
| KDR | X |
| EPHA5 | X |
| TET2 | X |
| INPP4B | X |
| FBXW7 | X |
| SDHA | X |
| TERT | X |
| IL7R | X |
| RICTOR | X |
| MAP3K1 | X |
| PIK3R1 | X |
| APC | X |
| RAD50 | X |
| CSF1R | X |
| PDGFRB | X |
| NPM1 | X |
| FGFR4 | see "Alterations Identified" |
| FLT4 | X |
| IRF4 | X |
| HIST1H3B | X |
| HIST1H1C | X |
| DAXX | X |
| PIM1 | X |
| CCND3 | X |
| PRDM1 | X |
| ROS1 | X |
| MYB | X |

| | |
|---|---|
| TNFAIP3 | X |
| ESR1 | X |
| ARID1B | see "Alterations Identified" |
| CARD11 | X |
| IKZF1 | X |
| EGFR | X |
| HGF | X |
| CDK6 | X |
| PIK3CG | X |
| MET | X |
| SMO | X |
| BRAF | X |
| EZH2 | X |
| KMT2C | X |
| FGFR1 | X |
| MYC | X |
| JAK2 | see "Alterations Identified" |
| CD274 | X |
| CDKN2A | X |
| CDKN2B | X |
| PAX5 | X |
| GNAQ | X |
| NTRK2 | X |
| FANCC | X |
| PTCH1 | X |
| ABL1 | X |
| TSC1 | X |
| NOTCH1 | see "Alterations Identified" |
| GATA3 | X |
| RET | X |
| JMJD1C | X |
| PTEN | X |
| SUFU | X |
| FGFR2 | X |
| HRAS | X |
| MYOD1 | X |
| WT1 | X |
| MEN1 | X |
| CCND1 | X |
| FGF19 | X |
| FGF4 | X |
| FGF3 | X |
| EED | X |
| MRE11 | X |
| ATM | X |
| SDHD | X |
| KMT2A | X |

| | |
|---|---|
| CBL | X |
| CHEK1 | X |
| KDM5A | X |
| CCND2 | X |
| ETV6 | X |
| CDKN1B | X |
| KRAS | X |
| ARID2 | X |
| ERBB3 | X |
| CDK4 | X |
| MDM2 | X |
| PTPN11 | X |
| HNF1A | X |
| CDK8 | X |
| FLT3 | X |
| FLT1 | X |
| BRCA2 | X |
| RB1 | X |
| IRS2 | X |
| NKX2-1 | X |
| TSHR | X |
| DICER1 | X |
| AKT1 | X |
| RAD51 | X |
| B2M | X |
| MAP2K1 | X |
| NTRK3 | X |
| IDH2 | X |
| BLM | X |
| IGF1R | see "Alterations Identified" |
| AXIN1 | X |
| TSC2 | see "Alterations Identified" |
| CREBBP | X |
| GRIN2A | X |
| SOCS1 | X |
| PALB2 | X |
| CBFB | X |
| CTCF | X |
| CDH1 | X |
| FANCA | X |
| TP53 | X |
| AURKB | X |
| MAP2K4 | X |
| FLCN | X |
| NF1 | X |
| SUZ12 | X |
| CDK12 | X |

| | |
|---|---|
| ERBB2 | X |
| RARA | X |
| BRCA1 | X |
| SPOP | X |
| RNF43 | X |
| PPM1D | X |
| BRIP1 | X |
| CD79B | X |
| PRKAR1A | X |
| RPTOR | X |
| SMAD2 | X |
| SMAD4 | X |
| BCL2 | X |
| STK11 | X |
| DOT1L | X |
| GNA11 | X |
| MAP2K2 | X |
| KEAP1 | X |
| SMARCA4 | X |
| BRD4 | X |
| JAK3 | X |
| PIK3R2 | X |
| MEF2B | X |
| CCNE1 | X |
| AKT2 | X |
| AXL | X |
| CIC | X |
| PPP2R1A | X |
| ASXL1 | X |
| SRC | X |
| TOP1 | X |
| AURKA | X |
| GNAS | X |
| RUNX1 | X |
| ERG | X |
| U2AF1 | X |
| CRKL | X |
| MAPK1 | X |
| SMARCB1 | X |
| CHEK2 | X |
| NF2 | X |
| EP300 | X |
| CRLF2 | X |
| BCOR | X |
| KDM6A | X |
| ARAF | X |
| GATA1 | X |

| | |
|---|---|
| KDM5C | X |
| AMER1 | X |
| AR | X |
| MED12 | X |
| ATRX | X |
| STAG2 | see "Alterations Identified" |
| BCORL1 | X |

Supplemental Figure 1.

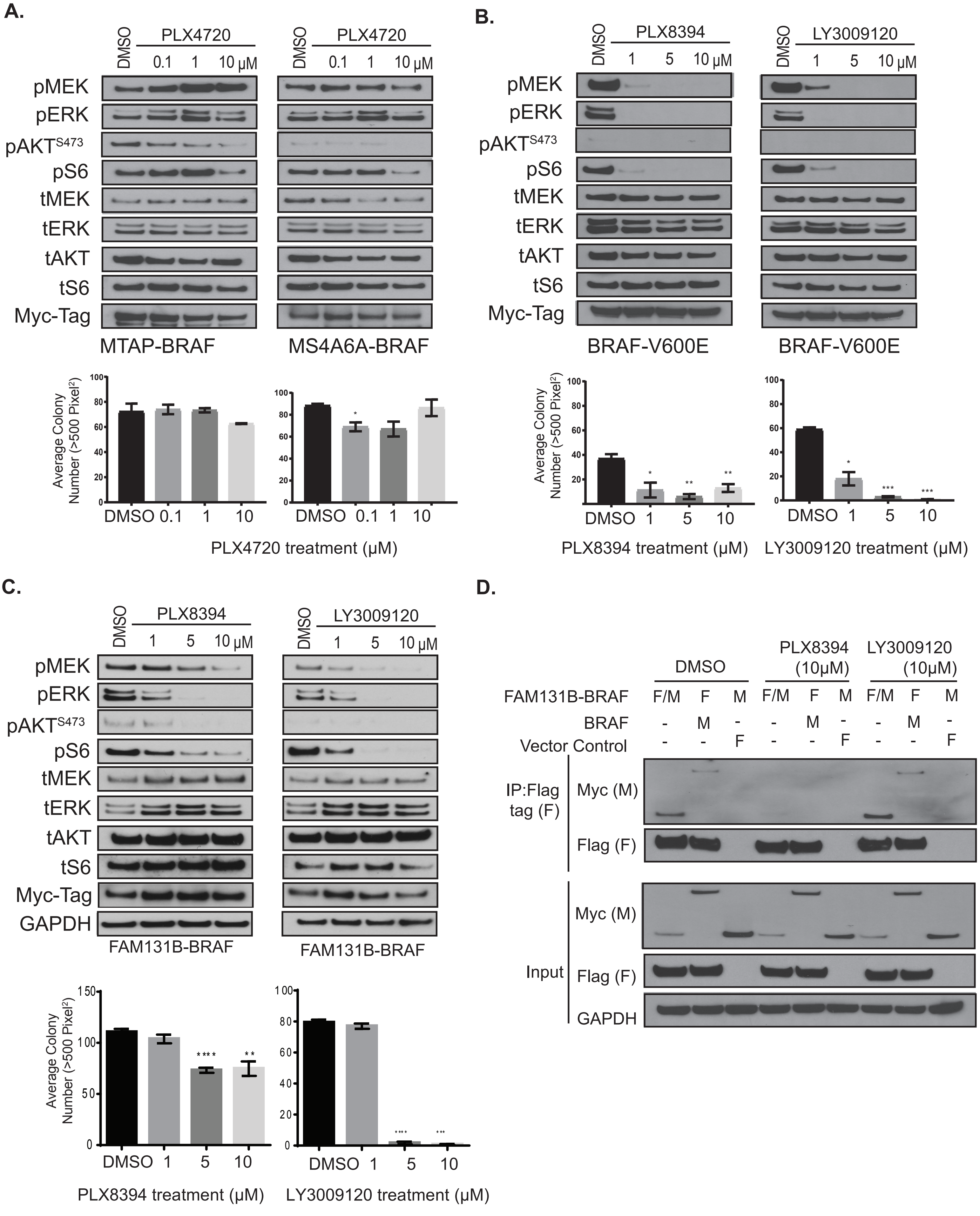

Supplemental Figure 2.

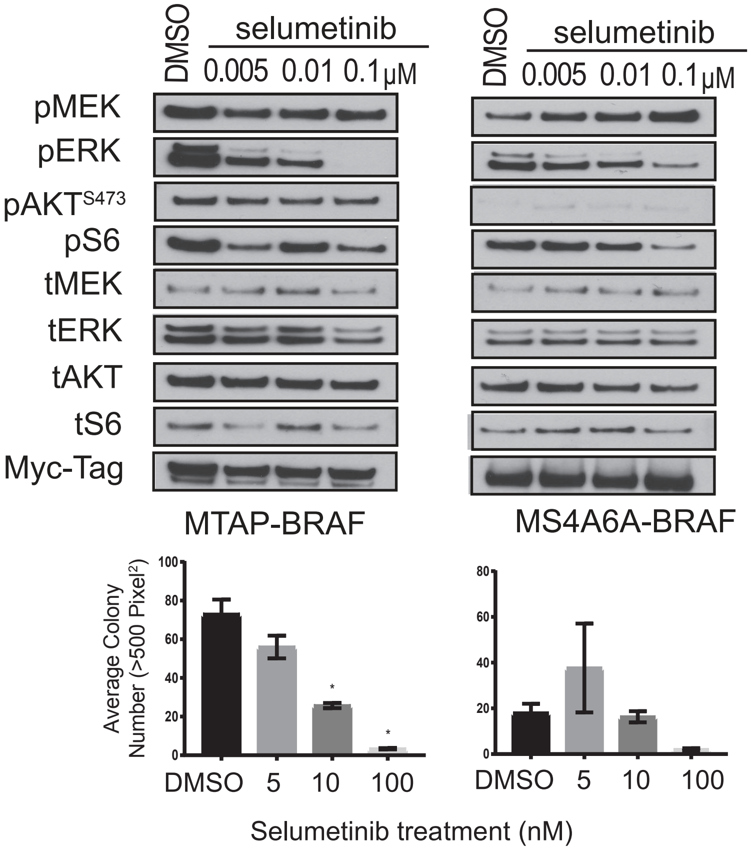
